# Tuning electrical spiking of *Schizophyllum* commune with light

**DOI:** 10.1101/2023.09.26.559269

**Authors:** Davin Browner, Andrew Adamatzky, Ella Schunselaar, Han A. B. Wösten, Phil Ayres

## Abstract

When studying the split-gill fungus *Schizophyllum commune*, we observed that the growing colonies displayed endogenous spikes of electrical potential similar to the action potentials of neurons. In order to investigate the impact of light on the electrical activities of these colonies, we exposed them to intermittent stimulation with cold light (5800k) and later with blue (c. 470nm), red (c. 642nm) and green (c. 538nm) light. Our findings revealed spiking activity can be influenced using this input including observable responses with patterns of spiking at relatively high average amplitudes (*>*1mV) appearing consistently upon illumination of the sample. The response is likely related to the activity of fungal photoreceptors, including potential sensitisation to blue light in the cellular signalling pathways facilitated by white collar proteins (WC-1, WC-2) in *S. commune*. Based on these findings, we suggest that fungal photosensors and photonic computing substrates have the potential to enable applications beyond the scope of conventional electronics via relatively fast spiking responses to light tuned by external input stimulation. Further work should focus on identifying the signal transduction pathway for responses to different wavelengths of light and its role in translation into engineered ELMs to extend existing studies in fungal photobiology.

## 1. Introduction

Living creatures generate electromagnetic fields and utilise them to acquire and process information about their environment, establish internal communication, and regulate their physical structure [1, 2, 3, 4, 5, 6, 7, 8, 9, 10]. From an information processing perspective, one of the most intriguing phenomena in bio-electricity is neural spiking. Spikes of electrical potential are well-known characteristics of neurons and are attributed to their learning and decision-making processes [11, 12, 13]. However, it is not just neurons that can generate spikes of electrical potential. Other living organisms, such as Protozoa [14, 15, 16], Hyrdoroza [17], slime molds [18, 19], and plants [20, 21, 22], also exhibit this capability. The occurrence of action potential-like spiking activity in fungi was first documented in 1976 [23], further confirmed in 1995 [24], and techniques for recording the electrical activity in fruiting bodies and colonized substrates were identified in 2018 [25]. In our efforts to understand the mechanisms of integrative electrical communication in fungi, we recorded and analyzed the electrical activity of various fungal species including oyster fungi *Pleurotus djamor* [25], bracket fungi *Ganoderma resinaceum* [26], ghost fungi (*Omphalotus nidiformis*), Enoki fungi (*Flammulina velutipes*), split gill fungi (*S. commune*), and caterpillar fungi (*Cordyceps militaris*) [27]. Our findings revealed significant variability in electrical spiking characteristics among these species, along with a complex nature of electrical communication events [28].

Light plays an important role in the growth and development of fungi. Fungi use numerous signal-transduction systems including photo-sensing being able to sense near-ultraviolet, blue, green, red and far-red light using up to 11 photoreceptors and signalling cascades to control a large proportion of the genome and thereby adapt to environmental conditions [29]. The photobiology of the ascomycete *Neurospora crassa* has received the most research attention including the effect of blue light on formation of sexual fruiting bodies [30] and the entrainment (resetting) of the circadian clock [31]. Blue light is detected by the white collar (WC) complex, which consists of the White Collar 1 and White Collar 2 (WC-1 and WC-2) proteins [32]. PAS (Per–Arnt–Sim) domains in WC-1 and WC-2 mediate the interaction between these proteins. The Light, Oxygen, Voltage (LOV) domain of WC-1 binds a flavin chromophore FAD (flavin adenine dinucleotide), and as such the complex can detect light. Upon light activation, the WC complex induces transcription of genes by binding to their promoter via the zinc finger DNA binding domains of WC-1 and WC-2. The light induced gene expression by the WC complex is transient. Light- exposed WC-1 becomes phosphorylated, which leads to inactivation of the complex and subsequent degradation of WC-1 [32]. Phytochromes, cryptochromes and rhodopsins are three other classes of photoreceptors [29]. Phytochromes detect red and far red light and are found in fungi, plants and bacteria. In plants, cryptochromes are blue light receptors and they show high sequence similarity to DNA photolyases, which are involved in light-dependent DNA repair.

Different light requirements have been described for mushroom production in basidiomycetes. Fruiting in the button mushroom *Agaricus bisporus* seems to be independent of light [33]. In contrast, *Coprinopsis cinerea* and *S. commune* need light for proper development of fruiting bodies. For *Coprinopsis cinerea* light ranging from 320 nm to 525 nm (i.e. ultraviolet, violet, blue, cyan, green) was shown to be responsible for induction of fruiting body formation. Blue light (c. 450nm 495nm) is necessary for initiation of mushroom formation in *S. commune* and the sequenced genome of this basidiomycete contains homologues of the blue light receptor genes in WC-1 and WC-2 of *Neurospora crassa*. This was observed via the subsequent formation of hyphal aggregates and primordia after stimulation with light where the response was linear with blue light intensity up to around 1250 lux. Inactivation of either or both of these genes has previously been shown to result in a blind phenotype [32]. Other wavelengths of light in the ultraviolet to red region are expected to have signal transduction functions in *S. commune* due to adjustment to different optical environments in wild samples. Comparative similarity to the light sensitive genes and proteins of the more extensively studied *Neurospora crassa* also support the likelihood that such a wavelength region would elicit signal transduction correlating to different behavioural functions.

The correlated optoelectrical spiking patterns for WC related photo-sensing and other photoreceptors are less explored in the general literature. In this paper, the electrical spiking correlates for the optical signal transduction pathway in response to cold light in samples of *S. commune* are examined. Investigating this photonic tuning of electrical spiking is useful in development of further understanding of the relationship between signal transduction and diverse behaviours including formation of fruiting bodies and augmentation of circadian clocks. Further, utilisation of these genetic and/or signal transduction pathways could be beneficial for the development of fungal photosensors and optical computing methods based on engineered living materials (ELMs). Optical ELMs have the potential to enable applications beyond the scope of conventional electronics where substrates are sensitised to particular photonic stimuli with the aim of design of optoelectronic elements with a high degree of spatiotemporal resolution and different application possibilities compared to wholly electrical circuits [34].

## 2. Methods

The strain H4-8A of the split-gill fungus *S. commune* was cultivated on *S. commune* minimal medium (SCMM) supplemented with 1.5% agar for a duration of three days at 30°C in a dark environment [35]. To ensure the integrity of the fungal colonies, the Petri dishes containing the colonies were not opened during the experiments and recordings for each of the three colonies were conducted after a total of 7-8 days of growth. Openings were created in the lids of the Petri dishes using a hot needle, and electrodes were carefully inserted through these openings until they reached the bottom of the dishes (refer to Fig. 1a). For recording the electrical activity of the fungal colonies, pairs of sub-dermal needle electrodes coated with iridium on stainless steel (manufactured by Spes Medica S.r.l., Italy) were utilised. These electrodes were equipped with twisted cables and connected to an ADC-24 high-resolution data logger from Pico Technology (UK), featuring a 24-bit analog-to-digital converter. The data logger ensured noise-free resolution through galvanic isolation and software-selectable sample rates. The distance between the electrodes within each differential electrode pair was c. 10 mm. Electrodes were arranged radially outwards from the inoculation point with an extent of 4 individual electrodes or 2 differential pairs.

**Figure 1:**
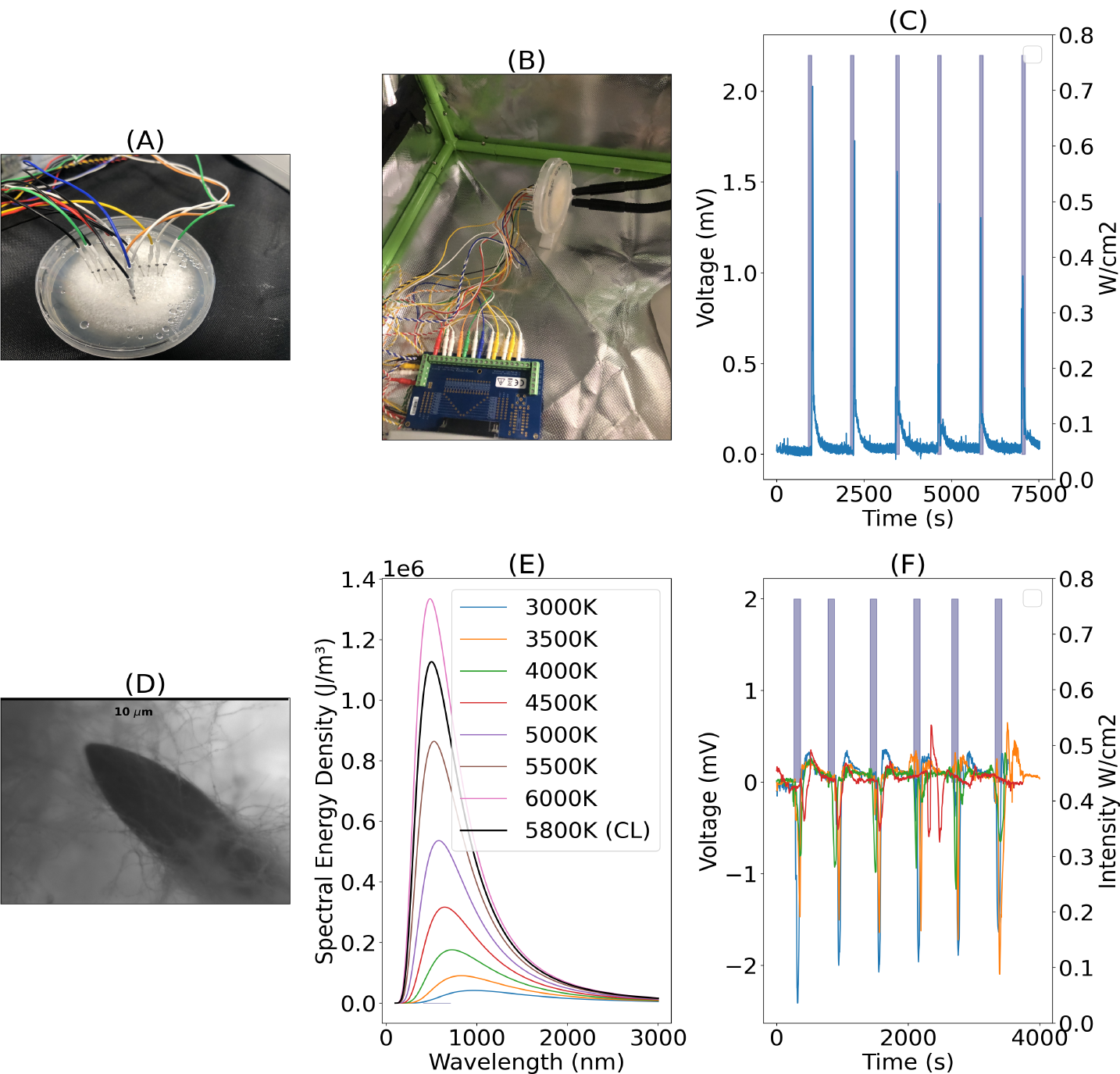
Setup and initial results of cold light illumination of *S. commune*. (a) The electrodes were arranged radially outwards from the centre of the Petri dish with an extent of four electrodes or two pairs of differential pairs of electrodes. (b) For all studies the Petri dish was fixed onto a stand with the underside pointed towards the light guide to occlude the electrodes from any direct optical interference or other similar effects. (c) In an exploratory temporal study the effect of cold light illumination was found to produce positive biased voltage spikes. Shaded regions indicate the duration of illumination events. (d) Growth of the fungal substrate onto the electrodes was imaged after two days in the grow tent showing hyphae connecting to the surface of the electrode. (e) Estimate of the wavelength and corresponding output of the cold light. (f) A further exploratory temporal study showed the effects of different illumination values (0.6 *W/cm*^2^, 0.5 *W/cm*^2^, 0.4 *W/cm*^2^, and 0.3 *W/cm*^2^ respectively) with lower intensity values producing lower corresponding mV output. With electrical spiking activity becoming The polarisation over time of the overall spiking response to a negative bias mV value was also observed.

The light modulation of the spiking frequency of the fungal substrate was investigated using the light source PHOTONIC PL-2000 (Ryf AG, Switzerland).

With halogen input and 0.6-0.7 maximum output or *W/cm*^2^ in intensity, the Photonic PL 2000 served as a constant cold white light source for stimulating the fungal substrate with light. The PL 2000 was calibrated to emit white light with a lamp voltage of 13 V. The lamp output optics were placed 4cm distance from the underside of the Petri dish containing the sample. A 3.5 mm light guide can produce a maximum of 0.8 W/cm2 of white light, whereas a 5 mm light guide can produce 0.6 W/cm2 of white light. The light source has minimal thermal output from the tip of the light guides and a temperature difference of 0.2-0.4^*°*^C (min-max reading over multiple cycles) between 0-0.6 *W/cm*^2^ (corresponding to c. 0-186 klux values) as measured on the surface of the Petri dish using a thermal camera (ETS320, FLIR, USA). For the colour filter study blue, green and red filters (Ryf AG, Switzerland) were added to the PL 2000. A compact CCD spectrometer (Thorlabs, USA) was used to characterise the wavelength and intensity of the optical inputs for the blue, green and red filters.

The electrical activity was recorded at a rate of one sample per second. Throughout the recording period, the data logger performed numerous measurements (typically up to 600 per second) and saved the average value. The acquisition voltage range was set at 78 mV. The recording was conducted continuously as the illumination was remotely activated without inducing mechanical vibrations in the sample, during which data was collected from 16 differential electrodes in 8 pairs. The electrodes were located on the opposite side of the Petri dish to the optical input. The inert coating of the electrodes prevented any interference with the growth of the fungal colonies, as depicted in Fig. 1b. During all experiments the fungal substrate was kept in the dark, with the exception of the illumination events, by placing the Petri dish and all recording equipment inside a light blocking tent.

## 3. Results

### 3.1. Temporal study

The initial exploratory temporal study revealed that the electrical spiking activity in *S. commune* could be tuned by cold light resulting in relatively faster inter-peak spike times as depicted in Fig. 1b compared to similar amplitudes (e.g. *>*1mV) observed in a previous study of multi-scalar endogenous spikes [36]. A linear relationship between the amplitude of intensity levels to amplitude in mV was observed. This pulsating behaviour in response to different light intensity values indicates that thresholds for spiking can be initiated. For example, no activity below a critical value and ability to toggle between two or more states based on the intensity of the photonic input and corresponding thresholds. Here, this is evident in Fig. 1f where indeterminant spiking responses were observed at values below 0.3 *W/cm*^2^ including delayed spikes at lower amplitudes. Progression of spiking activity to positive and negative electrical biases and overall polarisation of the electrode pairs was also observed but the cause was not immediately evident from this study. Onset of the optoelectrical spikes were observed to be relatively fast and more predictable compared to endogenous spiking (average of 400ms for maximum intensity values). This indicates that light stimulus latency of spiking is likely to be low and suitable for practical use such as implementation of threshold banded logic gates and in specific photosensing applications such as artificial photoreceptors.

### 3.2. Multi-trial temporal study

A second and more extensive temporal study was conducted where the fungal substrate response was recorded when exposed to multiple temporal illumination patterns of cold light over 10 trials of 10 stimulus events for each of the following ON/OFF patterns: 120s:240s, 120s:60s,60s:30s and 40s:20s as shown in Fig. 2. Here, polarisation of the signal over repeated illumination events was noted where early trials (1-5) positive bias mV values were identified with later trials (6-10) having prominent negative bias mV values. For a total of 10 *×* 10 trials (where there was 10 spikes for every trial) per temporal interval the number of spikes in each time period were automatically counted by utilising SciPy [37]. Specifically, the function FindPeaks was utilised to identify peaks with a prominence of 0.3 (here representing mV values) and distance value of 10. Peaks with a negative electrical bias were automatically counted using the same procedure but for negative bias values. This results in a basic method to dynamically tune the thresholds for electrical spiking activity even in the case where there is a gradual switch in polarity or a shift to positive or negative biases. Differences in spiking activity for electrode pairs that established connections with hyphae and those having indeterminate connections were observed. Over 10 trials per temporal interval the lack of signal transduction in the non-connected electrode pair can be clearly observed including in zoomed in portions of single trials. This is shown in Fig. 4 where the response of both connected and non-connected electrode pairs is depicted for the full recording encompassing all trials. Pronounced spiking activity is evident for the connected electrode pair corresponding to the illumination pattern whereas the non-connected electrode pair shows a typical noise output for agar. The zoomed in signal shows a single trial (1×10) and evidences the lack of periodic response for the agar electrode pair to the photonic input. Delayed onset of spiking (*>*1s), which was also observed in the initial exploratory studies, was observed for the first trial of ON:OFF 60s:120s but this can be seen as an anomaly due to consistent spiking onset following illumination for other trials.

**Figure 2:**
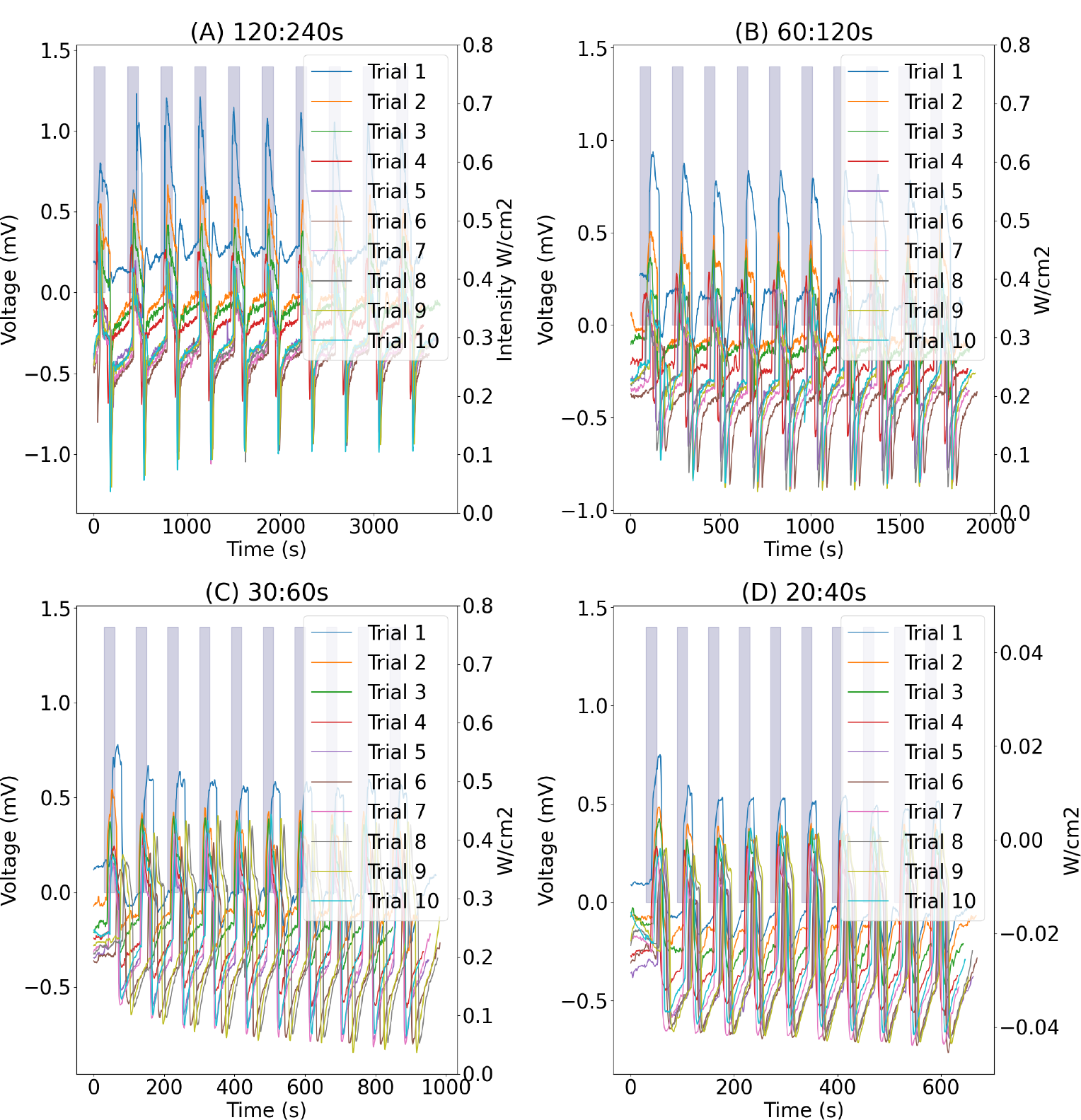
Temporal study of cold light illumination of *S. commune*.The fungal substrate response when exposed to multiple temporal illumination patterns of cold light including ON/OFF patterns of (a) 240s:120s, (b) 120s:60s, (c) 60s:30s and (d) 40s:20s. Shaded regions in each graph indicate the full period of illumination.

**Figure 3:**
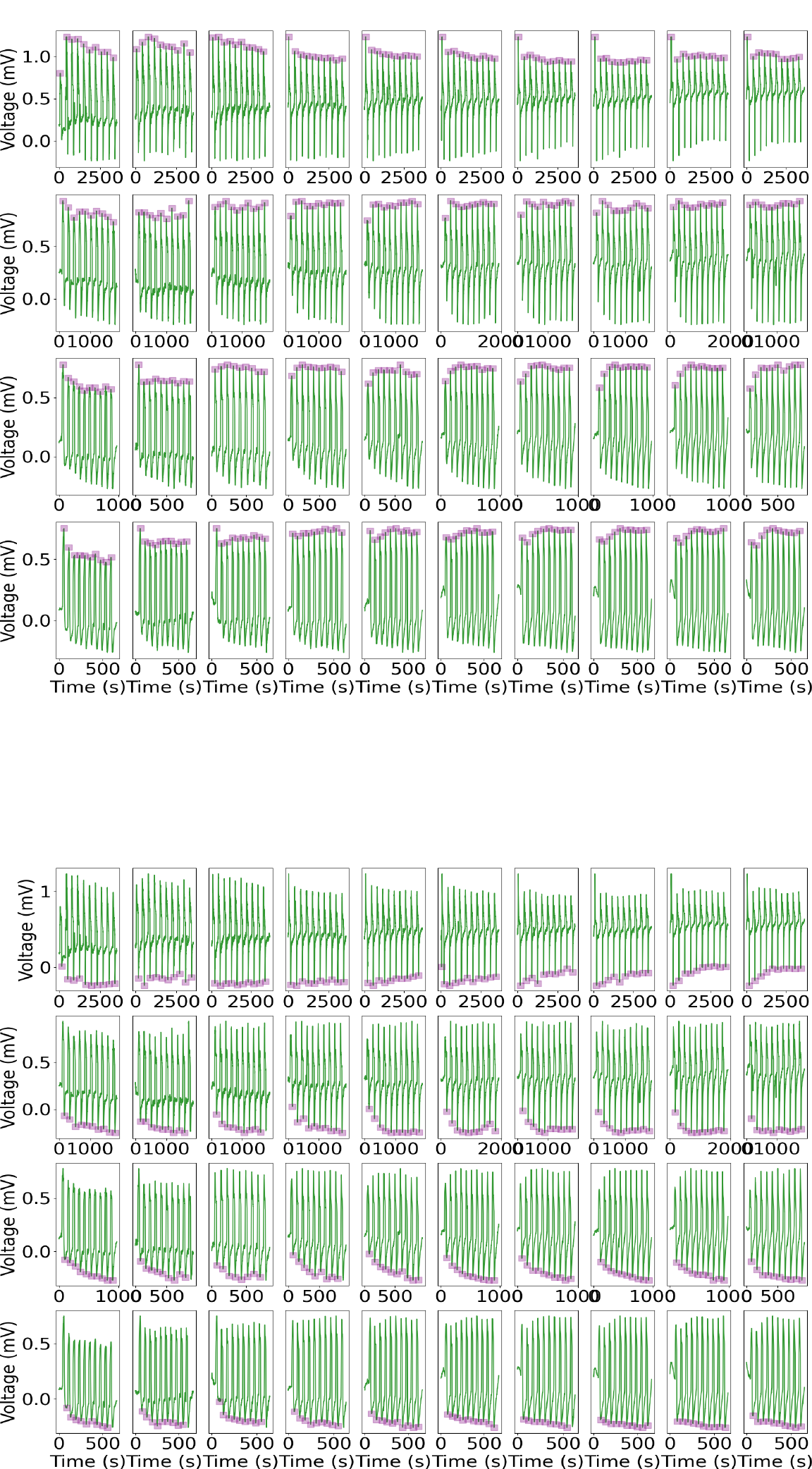
Peak detection. (a) positive bias voltage peaks (b) negative bias voltage peaks.

**Figure 4:**
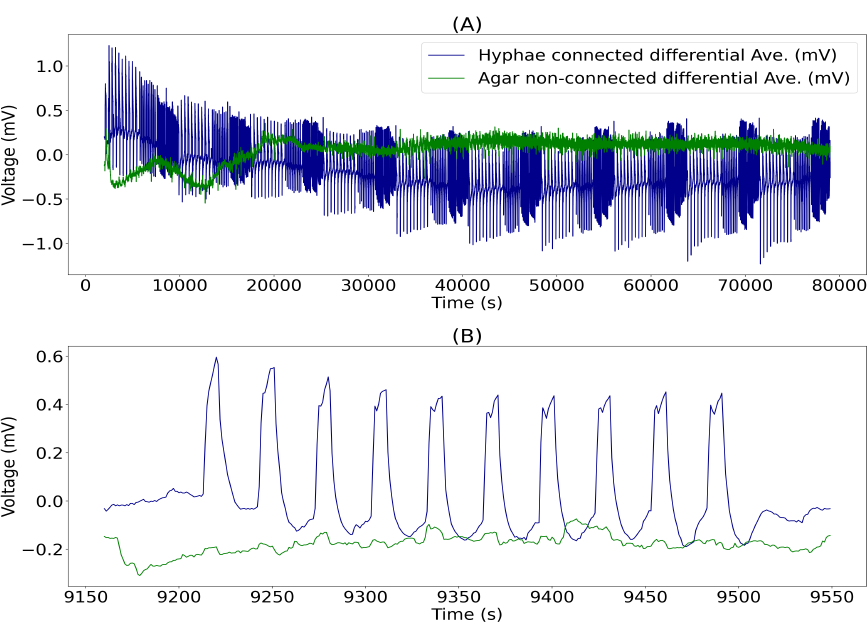
Signal vs. noise in electrodes verified via microscope imaging to be connected to hyphae and with no connection respectively. (a) Connected electrodes (dark blue) with optical responses over 10 trials of the temporal study contrasted with noise from non-connected electrodes with indeterminate connections to hyphae. (b) Zoomed in portion of graph (a) containing one trial during the temporal study and showing lack of response (i.e. no corresponding peaks) to photonic stimuli in the non-connected electrode versus the pronounced response to illumination in the connected electrode.

### 3.3. Identification of negative and positive biased peaks and intervals of spiking

The method for identification of amplitude peaks was used to estimate the inter-peak intervals for spikes and the distribution of amplitudes for both positive and negative biases (see Fig. 5). Progressive polarisation of the signal can be observed here as in Fig. 2. Average peak amplitudes recorded were calculated and are skewed to lower mean values as a result of drift and polarisation of the signal. For positive biased peaks: 0.31 mV for ON:OFF 120s:240s (median= 0.19mV, *σ*= 0.30mV), 0.30mV for ON:OFF 60s:120s (median= 0.21mV, *σ*= 0.2mV), 0.35mV for ON:OFF 30s:60s (median= 0.35mV,*σ*= 0.11mV), and 0.33mV for ON:OFF 40s:20s (median= 0.32mV,*σ*= 0.09mV). For minus amplitude peaks recorded values were: -0.75 mV for ON:OFF 120s:240s (median= -0.89mV,*σ*= 0.29mV), -0.64mV for ON:OFF 60s:120s (median= -0.77mV, *σ*= 0.24mV), -0.58mV for ON:OFF 30s:60s (median= -0.66mV,*σ*= 0.20mV), and 0.54mV for ON:OFF 40s:20s (median= -0.64mV,*σ*= 0.17mV). The average inter peak interval times for positive bias spikes were: 352.10s for ON:OFF 120s:240s (median= 360s,*σ*= 43.30s), 187.54s for ON:OFF 60s:120s (median= 180s,*σ*= 25.27s), 95.15s for ON:OFF 30s:60s (median= 90s,*σ*= 18.81s), and 64.93s for ON:OFF 40s:20s (median= 60s,*σ*= 17.75s). For negative biases similar intervals were apparent: 352.33s for ON:OFF 120s:240s (median= 360s,*σ*= 38.27s), 187.51s for ON:OFF 60s:120s (median= 180s,*σ*= 25.94s), 95.11s for ON:OFF 30s:60s (median= 90s,*σ*= 17.88s), and 64.77s for ON:OFF 40s:20s (median= 60s,*σ*= 17.95s). As calculated relative to local peaks (i.e. the maximum local value to the onset of the photonic signal) the average light stimulus latency of spiking for each temporal interval across all trials were: 67s for ON:OFF 120s:240s (median = 67.3s, *σ* = 29.4s), 29.22s for ON:OFF 60s:120s (median = 30.35s, *σ* = 18.5s), 14.76s for ON:OFF 30s:60 (median = 15.17s, *σ* = 8.94s), and 9.37s for ON:OFF 40s:20s (median = 9.35s, *σ* = 4.9s). Onset times (i.e. above noise floor response to light stimulus) averaged at <1s for all trials.

**Figure 5:**
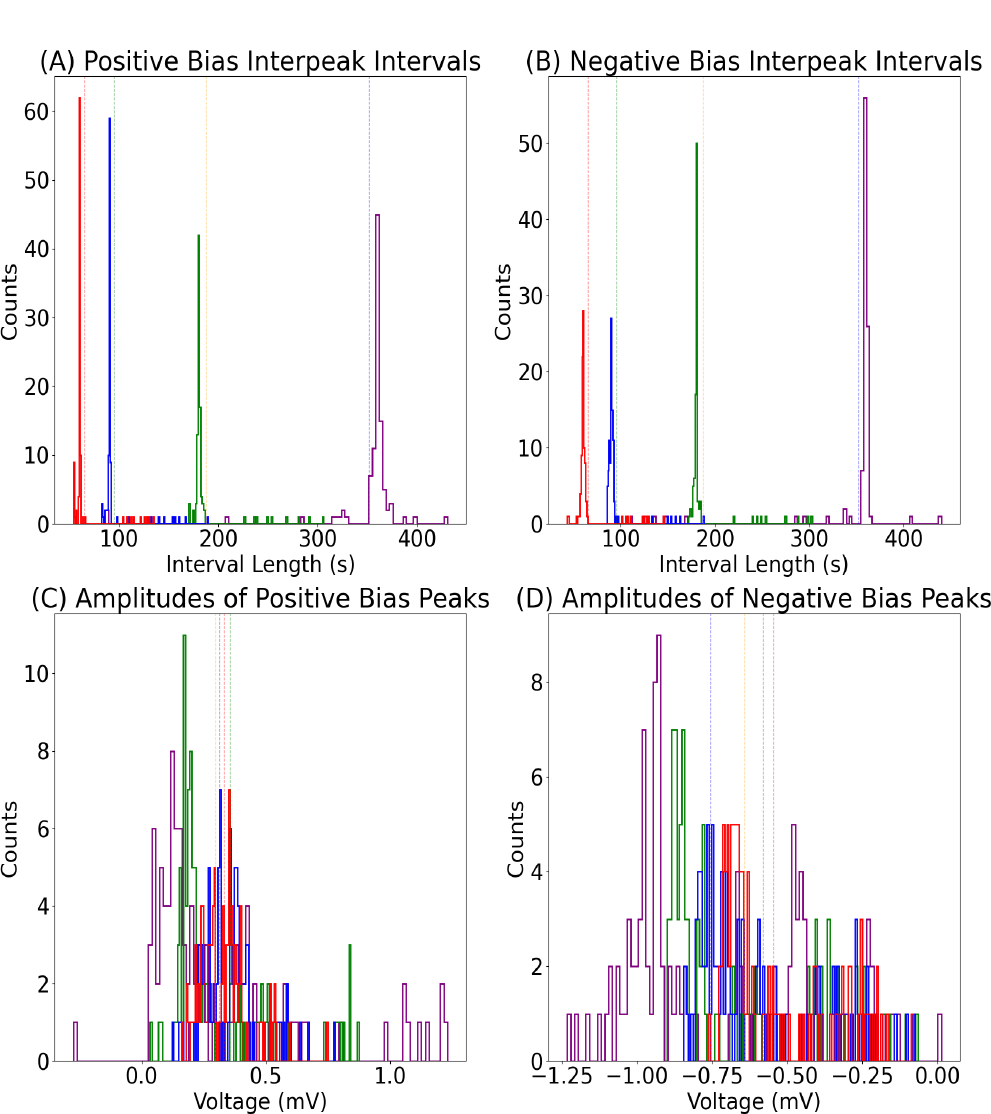
Histograms showing the inter-peak intervals and amplitudes of positive and negative bias voltage responses to cold light illumination of *S. commune*.(a) inter-peak intervals for positive bias voltage responses, (b) inter-peak intervals for negative bias voltage responses, (c) distribution of amplitudes for positive bias voltage responses and (d) distribution of amplitudes for negative bias voltage responses. For (a)-(d) mean values are indicated by dashed lines.

### 3.4. Repeat study and comparison with endogenous spiking

Another culture of *S. commune* was tested to give an overview of the potential for repetition of the results with identical photonic input. The repeat experiment showed similar electrical spiking responses with pronounced spiking on two pairs of differential electrodes as depicted in Fig. 6. For this repeat experiment the values peak amplitudes of spikes were lower (average of 0.5mV and 0.4mV for (a) and (b) respectively) compared to the previous results which showed more prominent repetition of positive and negative biased peaks of *>*1mV. For the 10×10 trials the latter culture did not show temporal polarisation of the signal forming an interesting difference with the previous result and indicating that stable thresholds can be set statically for implementation of logic and computation. In Fig. 6c the simultaneous recording of endogenous spiking patterns over a similar period can be observed for a section of the fungal substrate that produced no response to the photonic input. In Fig. 6d of the same figure a noise like response can be seen with some sporadic spiking.

**Figure 6:**
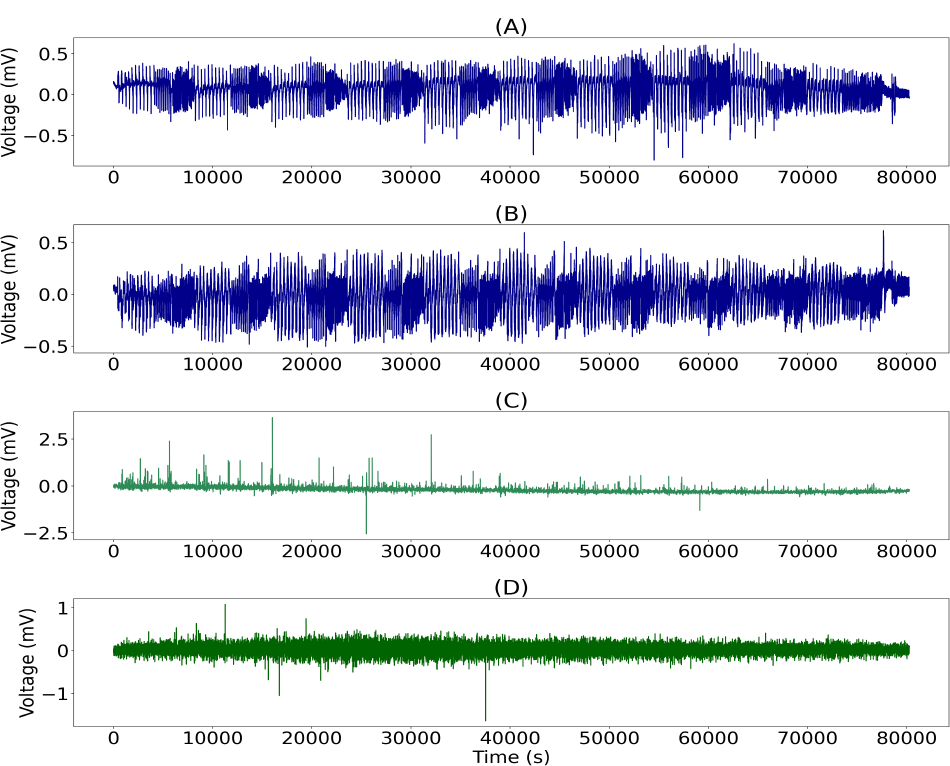
Repeat experiment on a second *S. commune* culture. (a) showing limited electrical polarisation (e.g. variation of spike response towards a negative bias) of the resulting signals. Indicating the possibility of implementation of stable thresholds for implementation of logic and computation,(b) simultaneous recording of a second differential channel showing similar response, (c) example of endogenous spiking over a similar time frame with positive bias spiking (*>*1mV) (d) an example of a more noisy recording on an indeterminately connected electrode.

### 3.5. Frequency and power spectrum analysis

The frequency and power spectrum characteristics of the resulting signals were also analysed as shown in Fig. 7. The average frequencies for each temporal study were: 0.029Hz for ON:OFF 120s:240s, 0.058Hz for ON:OFF 60s:120s, 0.10Hz for ON:OFF 30s:60, and 0.16Hz for ON:OFF 40s:20s. The power spectrum density (PSD) was estimated by computing the Welch PSD of each signal using the scipy.signal library [37]. First, the signal was divided into overlapping segments (using the Hanning window) and the periodogram of each segment was calculated with the periodiograms then averaged. The resulting average PSD for the overall signal across all trials for 120s:240s was 0.008 V^2^/Hz, 60s:120s was 0.010 V^2^/Hz, 30s:60s was 0.014 V^2^/Hz, and 20s:40s was 0.016 V^2^/Hz. The frequency axis and the PSD estimate of the photonic inputs can be seen in Fig. 7. These results indicate that the frequency of the electrical activity can be controlled via external optical stimuli.

**Figure 7:**
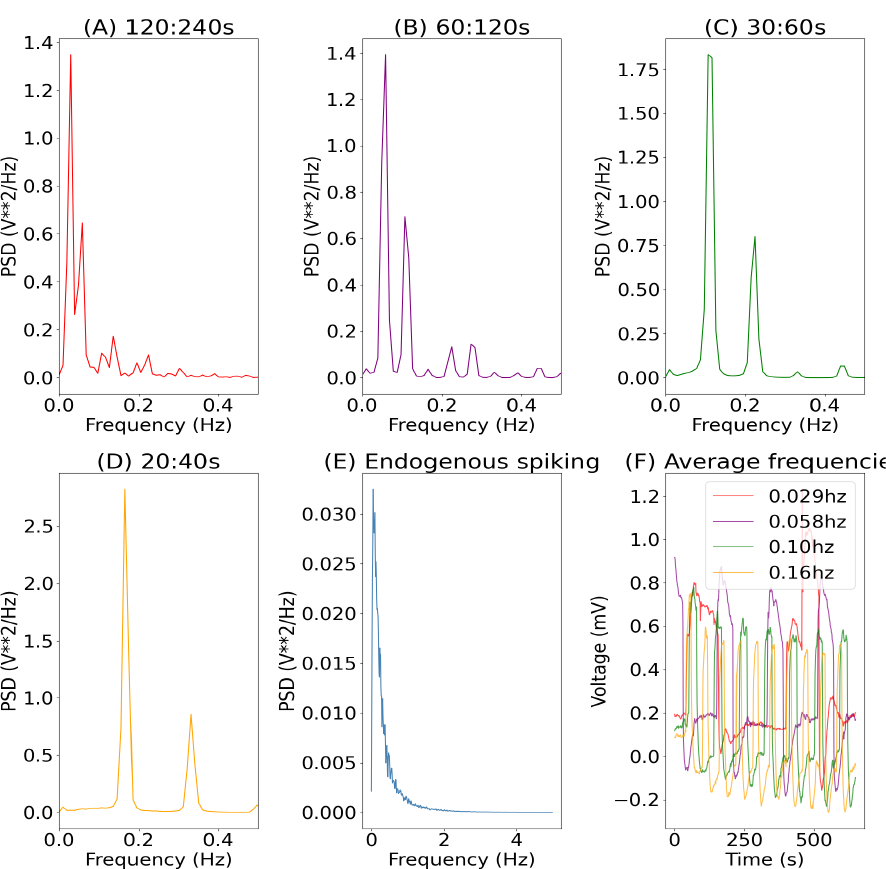
Frequency vs power spectrum comparison.(a) 240s:120s, (b) 120s:60s, (c) 60s:30s, (d) 40s:20s, (e) endogenous (no stimulation), (f) multi-frequency photonic spiking with corresponding average frequency values for the time period of recording.

### 3.6. Centrifugal waves of activity with chiral electrical polarisation

Some further tests revealed that the appearance of polarising positive and negative biases in response to the photonic input may be related to the spatial positioning of the electrodes where both electrodes made a suitable connection with the substrate hyphae. As depicted in Fig. 8c for testing of production of chiral waves of activity the electrodes were positioned at the furthest extent. As waves of electrical activity are centrifugal due to the way the mycelium in *S. commune* grows outwards from the inoculation point parallel electrode pairs to the radius should have pronounced spikes. Equally if the pair is perpendicular to the radius, the spikes may not be visible or particularly pronounced. In the case of our recordings of chiral waves, the wave was initiated between two electrode pairs that were parallel. This resulted in respective polarisation of the differential electrical potential as shown in Fig. 8a/b.

**Figure 8:**
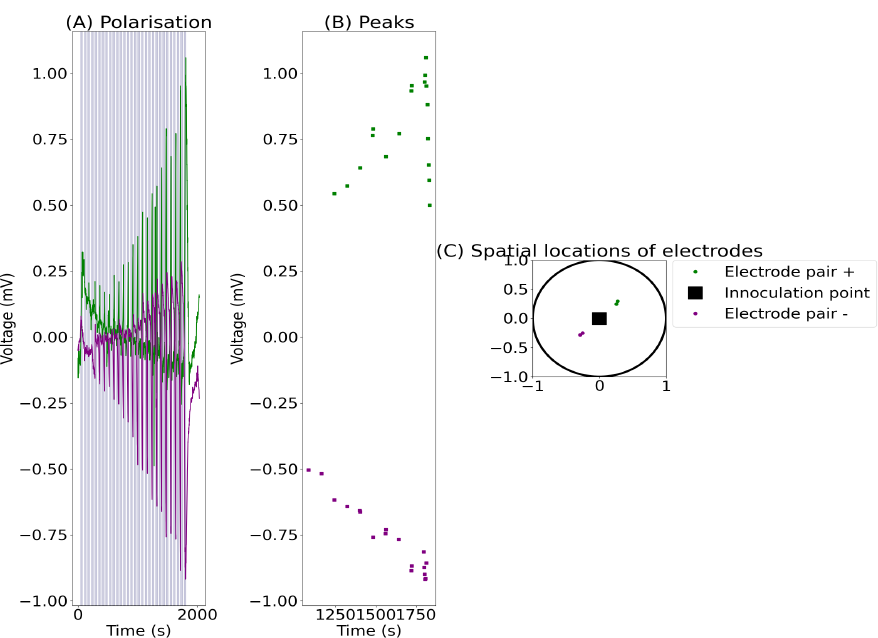
Chiral pattern in the electrical signal polarisation of recordings of two outermost electrode pairs exposed to repeated illumination (0.6 *W/cm*^2^ intensity).(a) two channels and illumination events (indicated by shaded regions) with positive and negative biases, (b) positive and negative bias peaks, (c) spatial position of electrodes.

**Figure 9:**
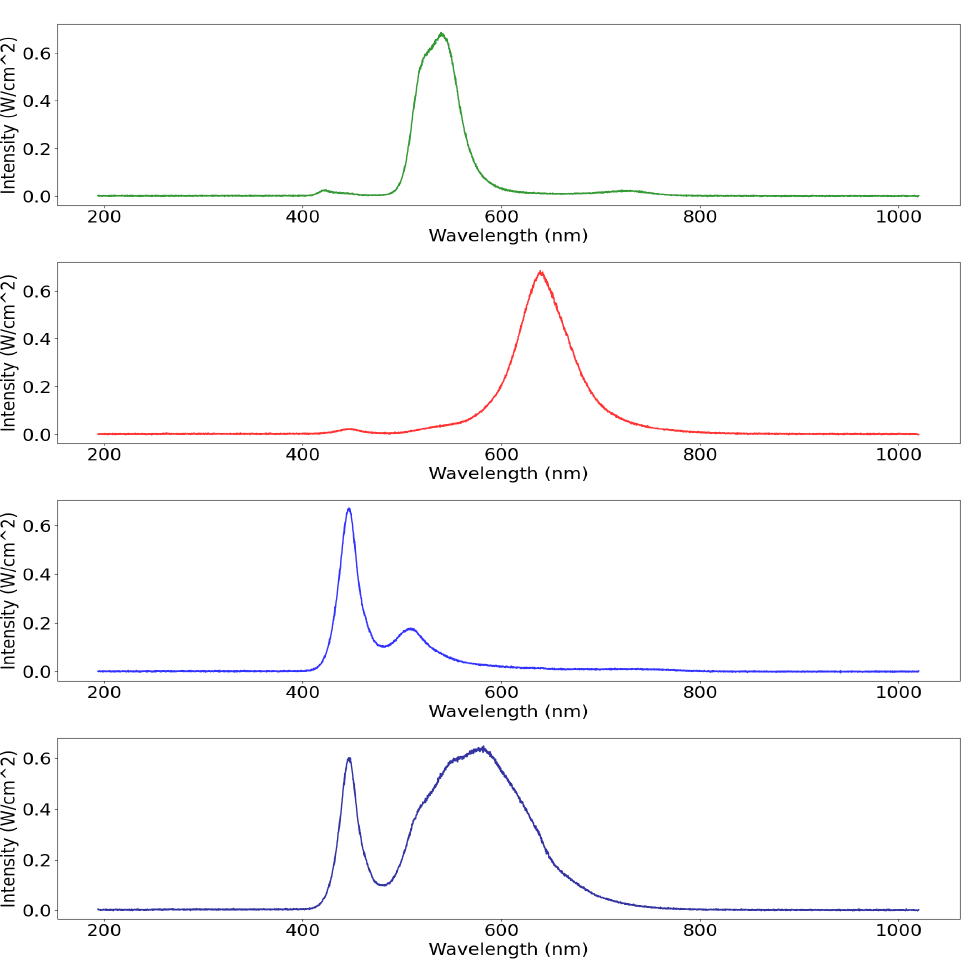
Spectral characterisation for (a) Blue, (b) Green (c) Red, (d) Cold light.

**Figure 10:**
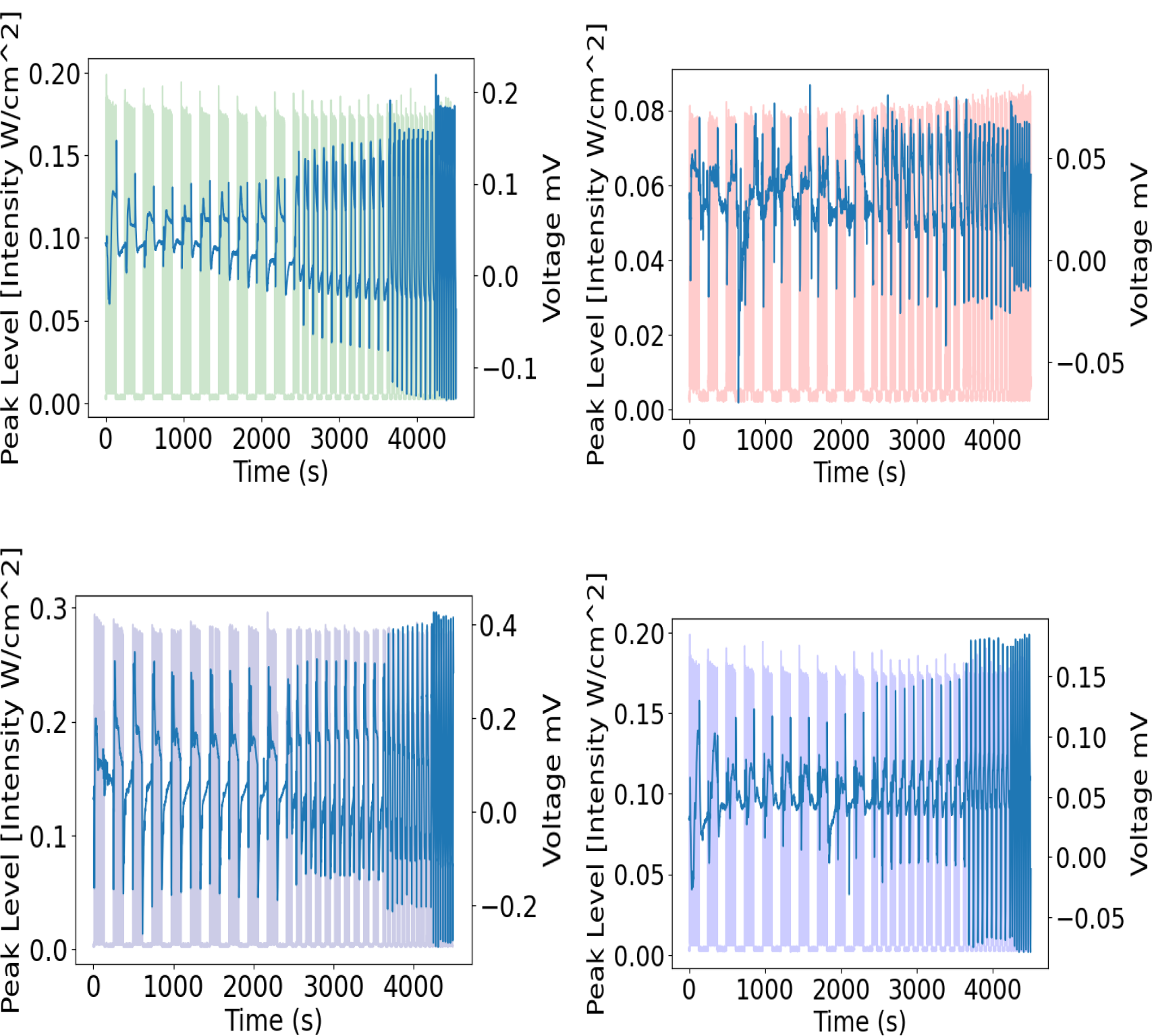
Illumination pattern and peak detection. Clockwise from top left: (a) Green, (b) Red, (c) Cold white light, (d) Blue.

**Figure 11:**
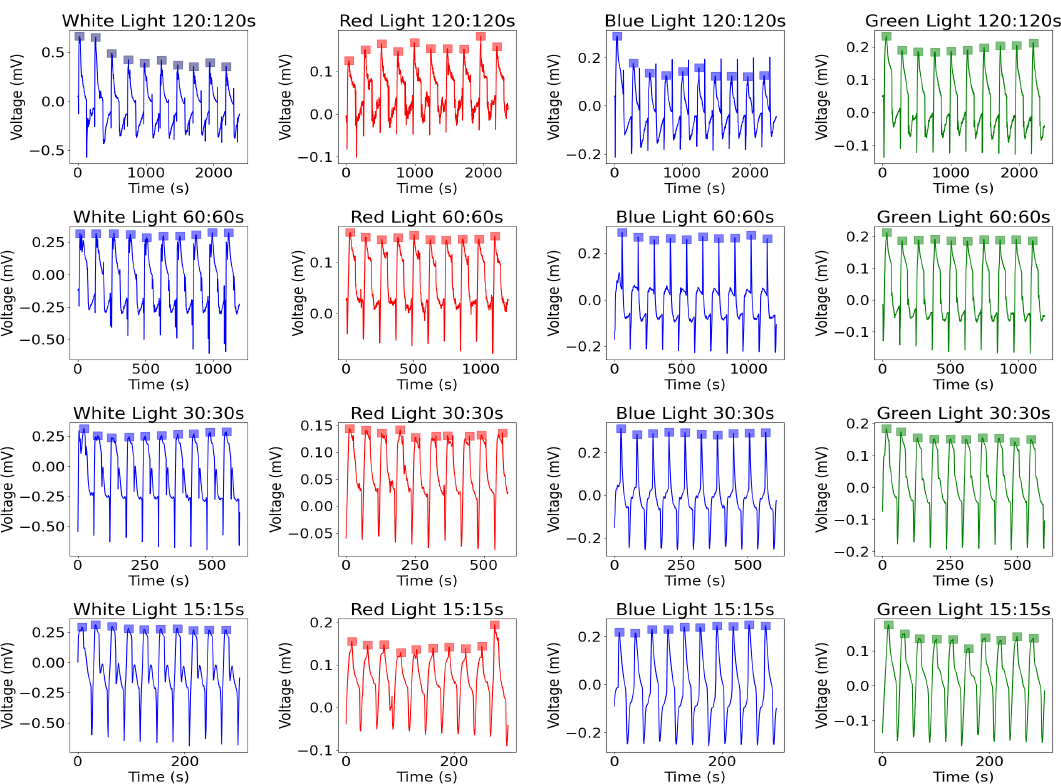
Peak detection for different stimulus time scales and different colours of light.

## 4. Wavelength and colour filter study

To explore specific colours and corresponding wavelength range of the light source for activation we repeated the experiment and stimulus parameters with the addition of blue, green and red colour filters. These patterns were compared to an identical cold/white light input pattern over a similar time frame. The mean weighted average wavelengths and intensities for each colour filter used in these experiments can be seen in Table 1.

**Table 1:**
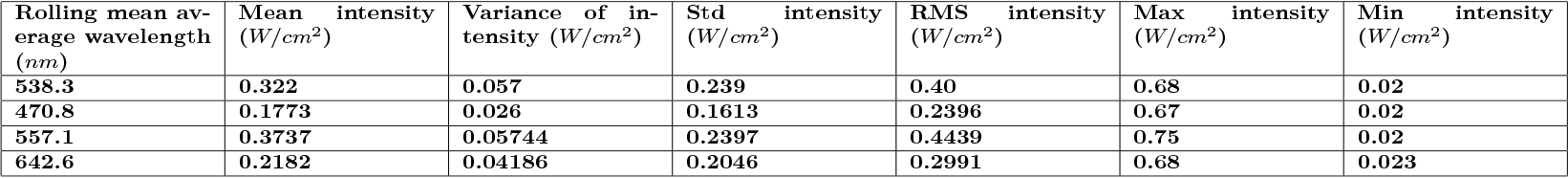
Characterisation of wavelengths and intensities of light source and filters.

| Rolling mean average wavelength<br>(nm) | Mean intensity<br>(W/cm<sup>2</sup>) | Variance of intensity<br>(W/cm<sup>2</sup>) | Std intensity<br>(W/cm<sup>2</sup>) | RMS intensity<br>(W/cm<sup>2</sup>) | Max intensity<br>(W/cm<sup>2</sup>) | Min intensity<br>(W/cm<sup>2</sup>) |
| --- | --- | --- | --- | --- | --- | --- |
| 538.3 | 0.322 | 0.057 | 0.239 | 0.40 | 0.68 | 0.02 |
| 470.8 | 0.1773 | 0.026 | 0.1613 | 0.2396 | 0.67 | 0.02 |
| 557.1 | 0.3737 | 0.05744 | 0.2397 | 0.4439 | 0.75 | 0.02 |
| 642.6 | 0.2182 | 0.04186 | 0.2046 | 0.2991 | 0.68 | 0.023 |

### 4.1. Spike latency of light stimulus

Intensity values were calculated relative to the highest intensity peak wavelength. As before the average light stimulus latency of spiking was calculated relative to local peaks (i.e. the maximum local value). Typical average values for the longest interval (120s:120s 0N:OFF) of spiking were: 22.8s (median = 21.5s, *σ* = 3.6s) for white light. Blue filtered light average was 46.1s (median = 48.0s, *σ* = 3.72s). Green filtered light average was 44.8s (median = 44.0s, *σ* = 4.51s). Red filtered light average was 36.5s (median = 36.0s, *σ* = 3.23s). For the interval of 60s:60s ON:OFF white light average was 26.6s (median = 28.0s, *σ* = 6.29s). Blue filtered light average was 57.9s (median = 58.0s, *σ* = 0.53 s). Green filtered light average was 25.2s (median = 25.0s, *σ* = 1.98s). Red filtered light average was 25.5s (median = 25.0s, *σ* = 1.68s). For shorter stimulus intervals including 30s:30s ON:0FF the average values for white light were 11.7s (median = 11.0s, *σ* = 3.97s). Blue filtered light average was 24.4 (median = 24.0s, *σ* = 0.48s). Green filtered light 9.9s (median = 10.0s, *σ* = 1.51s). For red filtered light the average was 15.7s (median = 12.0s, *σ* = 7.4s). Finally, for 15s:15s ON:OFF the average for white light was 4.4s (median = 4.5s, *σ* = 1.42s). Blue filtered light average was 8.7s (median = 9.0s, *σ* = 0.45s). Green filtered light average was 10.3s (median = 10.5s, *σ* = 0.9s). Red filtered light average was 8.9s (median = 9.5s, *σ* = 2.02s). Onset of spiking times (i.e. when the electrical potential passed above a minimum mV threshold distinct from the noise floor) averaged below 1s.

## 5. Discussion

Previous studies showed the importance of the role of photosensing in *S. commune* in relation to diverse behaviours including sexual reproduction [30] and regulation of circadian clocks [31]. The relationship between these signal transduction pathways in fungi and potential correlates in the form of electrical spiking have not been investigated previously. Here, we showed that there is a pronounced electrical activity in response to cold light (in this case 5,800K based on the output of the Photonic PL 2000). Cold light includes a significant amount of blue light (c. 450nm) which is one likely source of the activation of the photo signal transduction method as a result of the presence of WC-1 and WC-2 proteins [32]. The relationship between the spectral composition of the cold light was then pursued using filtered blue, green and red wavelengths indicating electrical spiking activity in response to all tested wavelengths although with differing prominences and widths in the resulting waveforms for each colour. Blue light was found to produce the highest amplitude of spikes when compared with green and red light. Further work is needed to identify specific spectral ranges and energy density of photonic inputs that activate this transduction pathway and correlated spiking activity including normalisation of intensity values using non-filtered (i.e. non-subtractive light input). More detailed experimental identification of the specific wavelengths and intensities that produce spiking activity would be useful. Either related to WC-1/WC-2 within a range close to blue light (450nm-495nm) or potentially for other light sensitive proteins at wavelengths including below 450nm including non-visible UV and over 490nm including IR ranges. The waveform of the input signals (e.g. sine, ramp, triangle, etc) should also be investigated as well as the type light source inputs (e.g. LEDs with intrinsic rather than filtered light inputs across different wavelengths). Later, when such a relationship is reasonably mapped the genetic basis for the production or non-production of spikes in response to blue light and other wavelengths can be augmented following the previous extensive work on light sensitive fungal proteins [32]. For example, allowing for production of patterned photosensors and computing substrates with many characteristics beyond the scope of conventional electronics including continuous regrowth of the substrate (e.g. as a result of damage) and synaptic photoreceptor output without the need for external circuitry such as multiple transistors (e.g. compared to a potential in-silico emulation using common circuit elements). In particular, in mycelial cultures where the intrinsic electrical dynamics of the substrate can be used to save energy and perform analog transduction methods that may be difficult to emulate in synthetic materials (e.g. highly spatially selective optoelectronic responses, optical initiation of chiral waves of electrical activity, etc). For computing, the presence of electrical spiking activity in response to all tested wavelengths of light suggests that logic gates may be implemented using the optoelectronic response in combination with static thresholds in mV values [38]. Comparatively, without digital implementation the demonstrated analog phototransduction method could be used to implement optical elements performing continuous time-domain computations locally [39, 40]. This would prove useful in the development of unconventional examples of analog optical computers using fungal substrates.

Considering fungal photobiology there are a large number of questions relating to the relationship between spikes and photonic input including a need for further elucidation of spiking responses to temporal and spatial variation of input signals, establishing the chemical basis for photosensing (e.g. experimentally mapping spiking patterns to specific light sensitive proteins), and identification of related amino acid sequences and translocation phenomena in mycelia. Only a small number of fungal photosensing genetic and signal transduction pathways have been identified in the literature and this research has been limited to a small number of fungi. The responses of *S. commune* to wavelengths other than blue light has not been properly studied in terms of correlated light sensitive proteins and genetic circuits. Single studies combining photoreceptor spike recordings and modification of light sensitive protein homologs (e.g. WC-1, WC-2, and deletions) would be useful for extending methods to study this signal transduction pathway. Such studies would allow for similar investigation of other wavelengths of light input producing spikes and corresponding proteins for the red and green spiking responses observed in this paper as well as other spectral regions. These methods can also be extended into studies of autoflourescence and bioluminescence in mycelium cultures and correlated electrical spiking activity. In terms of electrical activity, the related functions of different spiking events is currently not well established and their correlation to external stimuli. More fine grained experiments including using real time calcium imaging and similar approaches would allow for further advancement of knowledge about the functional correlates to the signal-transduction pathways excited by light. These studies can help to establish differences between specific physiological functions and potential epiphenomena such as photoelectric responses of fungal pigments which form part of the fungal skin and related spiking activity [41].

## 6. Acknowledgement

The research has been conducted under the framework of the FUNGATERIA (www.fungateria.eu) project, which has received funding from the European Union’s HORIZON-EIC-2021-PATHFINDER CHALLENGES programme under grant agreement No. 101071145. It is co-funded by the UK Research and Innovation grant No. 10048406.

## 7. Data availability

The datasets used and/or analysed during the current study available from the corresponding author on reasonable request.

